# Expanding vaginal microbiome pangenomes via a custom MIDAS database reveals *Lactobacillus crispatus* accessory genes associated with cervical dysplasia

**DOI:** 10.1101/2025.09.11.675634

**Authors:** Claire A. Dubin, Chunyu Zhao, Katherine S. Pollard, Tomiko Oskotsky, Jonathan L. Golob, Marina Sirota

## Abstract

The vaginal microbiome plays a central role in reproductive health. Vaginal microbiome dysbiosis is associated with many adverse reproductive health outcomes, but most studies have focused on associations at the species level. The potential contribution of intraspecies microbial variation, especially gene content differences across bacterial strains, remains underexplored in reproductive health contexts. The Metagenomic Intra-Species Diversity Analysis (MIDAS) framework enables such analyses but depends on comprehensive reference databases. We constructed a MIDAS-compatible pangenome database from over 18,000 genomes in the Vaginal Microbiome Genome Collection (VMGC). Compared to the Genome Taxonomy Database (GTDB)-derived reference, the VMGC database expanded pangenomes of prevalent vaginal species, better capturing vaginal-specific intraspecies diversity. Applying this database to vaginal samples from a cervical dysplasia cohort, we identified thirteen *Lactobacillus crispatus* accessory genes significantly associated with cervical dysplasia, including a HicAB toxin-antitoxin system, three transcriptional regulators, and three phage-derived genes. These findings highlight the utility of body site-specific reference resources and shotgun metagenomic sequencing for uncovering intraspecies microbial variation relevant to reproductive health.

**IMPORTANCE:** The vaginal microbiome plays a critical role in reproductive health, and different bacteria from the same species can carry different genes that influence how the strains interact with the host and other microbes. These strain-level differences are often overlooked when microbiomes are analyzed only at the species level. Existing genomic reference databases are heavily biased toward gut and environmental bacteria, leaving the genetic diversity of vaginal microbes understudied. We built a specialized reference database from over 18,000 vaginal bacterial genomes that better reflects this diversity. We then applied this resource to quantify gene-level variation in vaginal samples from a cervical dysplasia cohort. Focusing on *Lactobacillus crispatus*, a prevalent and often beneficial vaginal species, we identified thirteen genes that were more common in women with cervical dysplasia than in controls. This work demonstrates that body site-specific genomic resources are essential for uncovering strain-level bacterial differences relevant to reproductive health.

## INTRODUCTION

The microbiome is increasingly recognized as an essential factor in health and disease throughout the human body.^1^ While the gut is the most well-studied system in microbiome research, microbial communities in other body sites also play an important role in host physiology and disease susceptibility.^2,3^ Bidirectional interactions between the vaginal microbiome and host play a key role in reproductive health.^4^ Disruption to this balance, often called vaginal dysbiosis, is linked to adverse reproductive health outcomes, including preterm birth^5–7^ and endometriosis.^8,9^ Vaginal dysbiosis involves a shift away from *Lactobacillus*-dominated communities toward more diverse and potentially pathogenic taxa.^4^ Pathogenic bacteria in the vaginal microbiome compromise the epithelial barrier and weaken immune responses, heightening susceptibility to infection by other pathogenic bacteria, yeast, and viruses.^4,10^ For example, persistent human papillomavirus (HPV) infection is the primary precursor to cervical dysplasia,^11,12^ the precancerous growth of abnormal cells on the cervix. Cervical dysplasia is a significant public health concern due to its potential to progress to cervical cancer, the fourth most common cancer in women.^12,13^ Further clarifying the role of the vaginal microbiome in preventing or promoting cervical dysplasia is essential for elucidating its underlying etiology, identifying biomarkers, and developing therapeutics.

While the majority of vaginal microbiome studies have focused on composition at the species level, intraspecies genomic variation also plays a role in the functional potential and ecological dynamics of the vaginal microbiome. *Lactobacillus crispatus*, a widely prevalent species often regarded as a marker of a healthy vaginal microbiome, has a large accessory genome compared to other common vaginal *Lactobacillus* species.^14^ Frequent gain and loss of genes across *L. crispatus* strains results in genetic and phenotypic variation among members of the species. *L. crispatus* vaginal isolates vary in their ability to utilize glycogen sources, produce hydrogen peroxide, and survive acidic stress.^15^ Genes encoding mucin-binding proteins and bacteriocins, which enhance adhesion to host tissues and inhibit pathogen growth, respectively, are variably present across *L. crispatus* strains.^15–17^ Vaginal *L. crispatus* strains differ in their ability to suppress the growth of bacterial and fungal pathogens, and *L. crispatus* and other common vaginal microbiome species exhibit different co-occurrence patterns at the strain level.^18^ These observations suggest that *L. crispatus* may not universally inhibit progression to a dysbiotic vaginal microbiome state.^19^ Recognizing and characterizing this intraspecies variation is essential for identifying reliable biomarkers of disease and developing probiotics to support reproductive health.

The Metagenomic Intra-Species Diversity Analysis (MIDAS) pipeline is a versatile tool for measuring intraspecific variation at the nucleotide and gene level from shotgun metagenomic sequencing data.^20,21^ MIDAS can be used to quantify bacterial gene content in metagenomic samples by mapping metagenomic reads to a preconstructed pangenome reference database. Intraspecies gene presence/absence data generated by MIDAS can be used in metagenome-wide association studies (MWAS) to identify bacterial accessory genes associated with clinical outcomes. For instance, recent work using the MWAS-based model microSLAM identified a fructoselysine utilization operon in *Faecalibacterium prausnitzii* that is negatively associated with inflammatory bowel disease.^22^ Given the demonstrated utility of MIDAS and microSLAM in identifying accessory genes associated with disease states and the mounting evidence for intraspecies genotypic and phenotypic variation in the vaginal microbiome, we sought to optimize this framework to identify associations between accessory genes in common vaginal microbiome species and reproductive health.

In this work, we address the limited representation of vaginal-derived genomes in existing MIDAS pangenome databases by developing a pipeline for constructing MIDAS-compatible custom pangenome reference databases and applying it to construct a pangenome database from a collection of over 18,000 vaginal genomes. We demonstrate that this database expands the pangenomes of six prevalent vaginal microbiome species to better reflect the genetic diversity of the vaginal microbiome compared to existing MIDAS pangenome databases. We then apply the newly constructed database to quantify gene content in vaginal microbiome samples and identify thirteen *L. crispatus* accessory genes associated with cervical dysplasia, including a toxin-antitoxin system and phage-derived genes of unknown function.

## RESULTS

### Construction of a MIDAS-compatible pangenome reference database from vaginal genomes

Strain-level association analyses rely on comprehensive and representative reference genomes. Reference-based gene content analyses are dependent on the quality of the reference database; genes that are absent from the reference cannot be measured, and an excess of irrelevant genes can introduce unnecessary noise. We set out to develop a MIDAS-compatible pangenome database specifically for use with vaginal microbiome specimens to refine and expand the landscape of genes available for reproductive health analyses. Two prebuilt MIDAS-compatible databases were previously constructed from genomes in the Unified Human Gastrointestinal Genome collection (UHGG)^23^ and the Genome Taxonomy Database r202 (GTDB).^24^ UHGG exclusively contains genomes from the human gut microbiome, and although GTDB contains a broader set of genomes, genomes from gastrointestinal samples make up approximately 35% of human-derived GTDB genomes with specified origin. In contrast, genomes from other human body sites such as the female reproductive tract, oral cavity, and skin make up only 1%, 2%, and 4%, respectively. Because more than 80% of GTDB r202 genomes are derived from cultured isolates, reference databases built from these genomes may not fully capture the uncultured and strain-diverse taxa prevalent in these underrepresented body sites. Recent collections comprised primarily of metagenome-assembled genomes (MAGs) have increased the number of available genomes for species in these less-studied niches.^25–27^ Expanding pangenome databases to include these MAGs may better reflect the microbial genetic diversity within these body sites.

We constructed a custom MIDAS-compatible pangenome reference database using 19,542 prokaryotic genomes in the Vaginal Microbiome Genome Collection (VMGC).^25^ The vast majority of genomes in VMGC (95%) are custom-assembled MAGs. Only 3% of the genomes in the VMGC are also present in the existing GTDB-based MIDAS reference, and these shared genomes are all derived from isolate sequencing. The VMGC prokaryotic genome assemblies and their GTDB 214.1 taxonomic assignments were downloaded from Huang et al (2024).^25^ There is a median of 19 input genomes per species (IQR 7.0-93.5), and genome quality ranges from medium-quality (52%), to high-quality (43%), and near-complete (5%) as calculated in Huang et al. (2024).^25^ We developed a Nextflow pipeline to streamline construction of a MIDAS-compatible pangenome database from a user-specified set of genomes and their species assignments as described in Zhao et al. (2023)^21^ and Smith et al. (2025)^20^ (Figure 1A). Using this pipeline, we created a pangenome reference database for each prokaryotic species with at least five assigned genomes, totaling 18,970 genomes across 179 species, 81 genera, and 42 families. Predicted genes from all genomes assigned to a species were clustered by average nucleotide identity (ANI) thresholds ranging from 75-99%. Higher ANI thresholds can distinguish subtle differences among highly similar genes, while lower thresholds can group more distant homologs. To balance detection of gene variants and grouping of gene families, we focused our pangenome analysis on 90% ANI gene clusters. The number of 90% ANI gene clusters per species ranges from 713 to 18,533 (IQR 2358.0-4409.5) and is positively correlated with the number of input genomes (Figure 1B, Spearman ρ = 0.57, *p* = 5e-17, Table S1). This trend is consistent across all tested ANI thresholds, with pangenome size decreasing as gene clusters are merged at lower ANI thresholds (Figure S1, Table S2).

**Figure 1.**
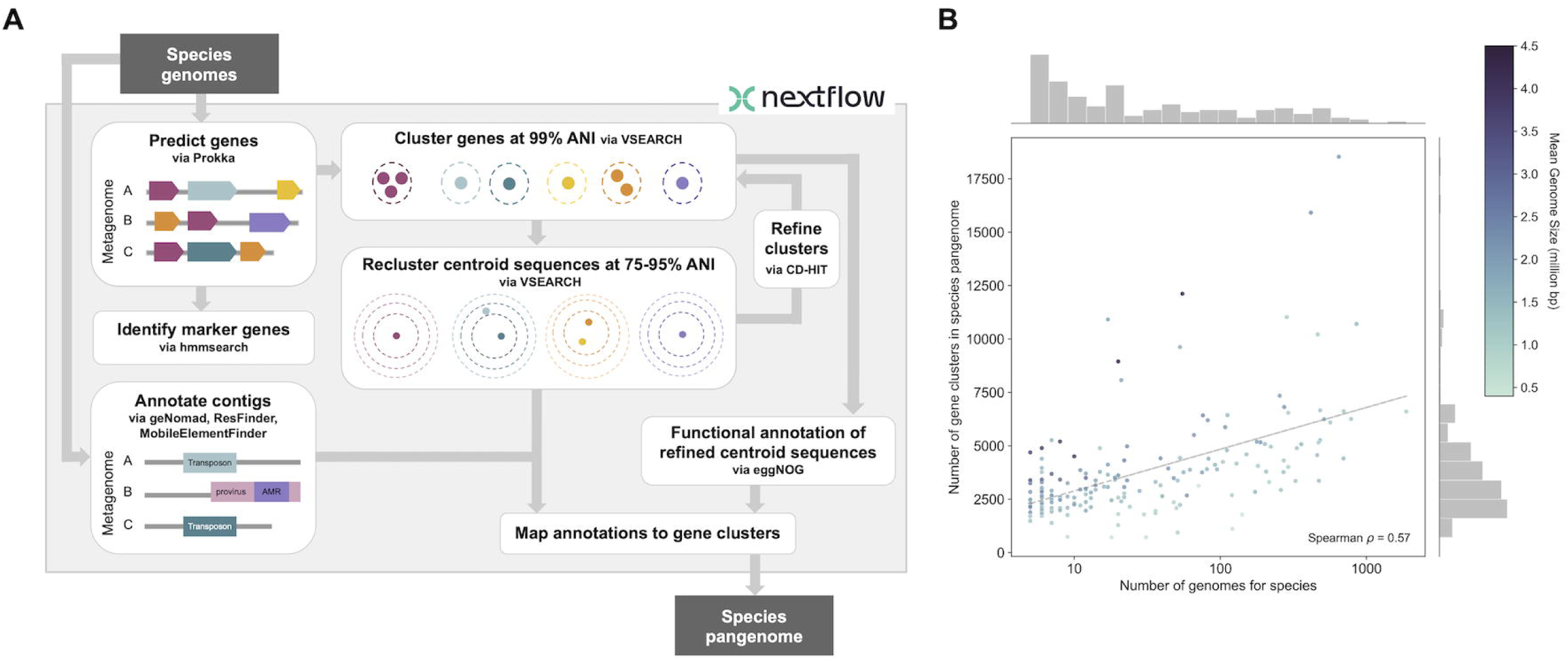
Construction of a custom pangenome database. **A.** Schematic for Nextflow pipeline for constructing a custom pangenome database from genomes of a given species. ANI = average nucleotide identity, AMR = antimicrobial resistance. **B.** Number of genomes vs. number of 90% ANI gene clusters for each species with at least 5 genomes in VMGC. Each point represents a species, and points are colored by average genome size for the species. Marginal histograms show the distributions of each variable: number of genomes per species (top) and number of 90% ANI gene clusters in species pangenome (right).

### Comparison to GTDB-only database

We next compared the ability of the VMGC-derived MIDAS pangenome reference database to reflect the genetic diversity of organisms in the vaginal microbiome as compared to the GTDB-derived reference.

Pangenome expansion widens the set of prokaryotic genes that can be tested for differential prevalence across strains and associated with clinical data. We compared pangenome sizes (number of 90% ANI gene clusters) of 129 species with one-to-one mappings across GTDB versions r202 and r214.1, which were used to determine taxonomic classification in the VMGC and GTDB databases, respectively (Table S1). Of these species, 66% (85/129) have a larger pangenome in the database constructed from VMGC genomes, and 34% (44/129) have a larger GTDB pangenome (Figure 2A). Differences in pangenome size are positively correlated with differences in the number of genomes per species in each database (Figure S2, Spearman ρ = 0.92, *p* = 8x10^-55^). The five species with the greatest difference in pangenome size between GTDB and VMGC, all with larger GTDB pangenomes, are gut-associated species (*Phocaeicola vulgatus, Enterococcus faecalis, Klebsiella pneumoniae*) and/or widely sampled pathogenic species (*Acinetobacter baumannii, Escherichia coli*). Species with a larger VMGC pangenome include all four hallmark *Lactobacillus* species used to classify vaginal community state types (*L. crispatus, L. gasseri, L. iners,* and *L. jensenii*),^28^ as well as species associated with vaginal microbiome dysbiosis, including six *Bifidobacterium vaginale* species (collectively referred to as *Gardnerella vaginalis* in NCBI taxonomy) and *Fannyhessea vaginae* (formerly named *Atopobium vaginae*).

**Figure 2.**
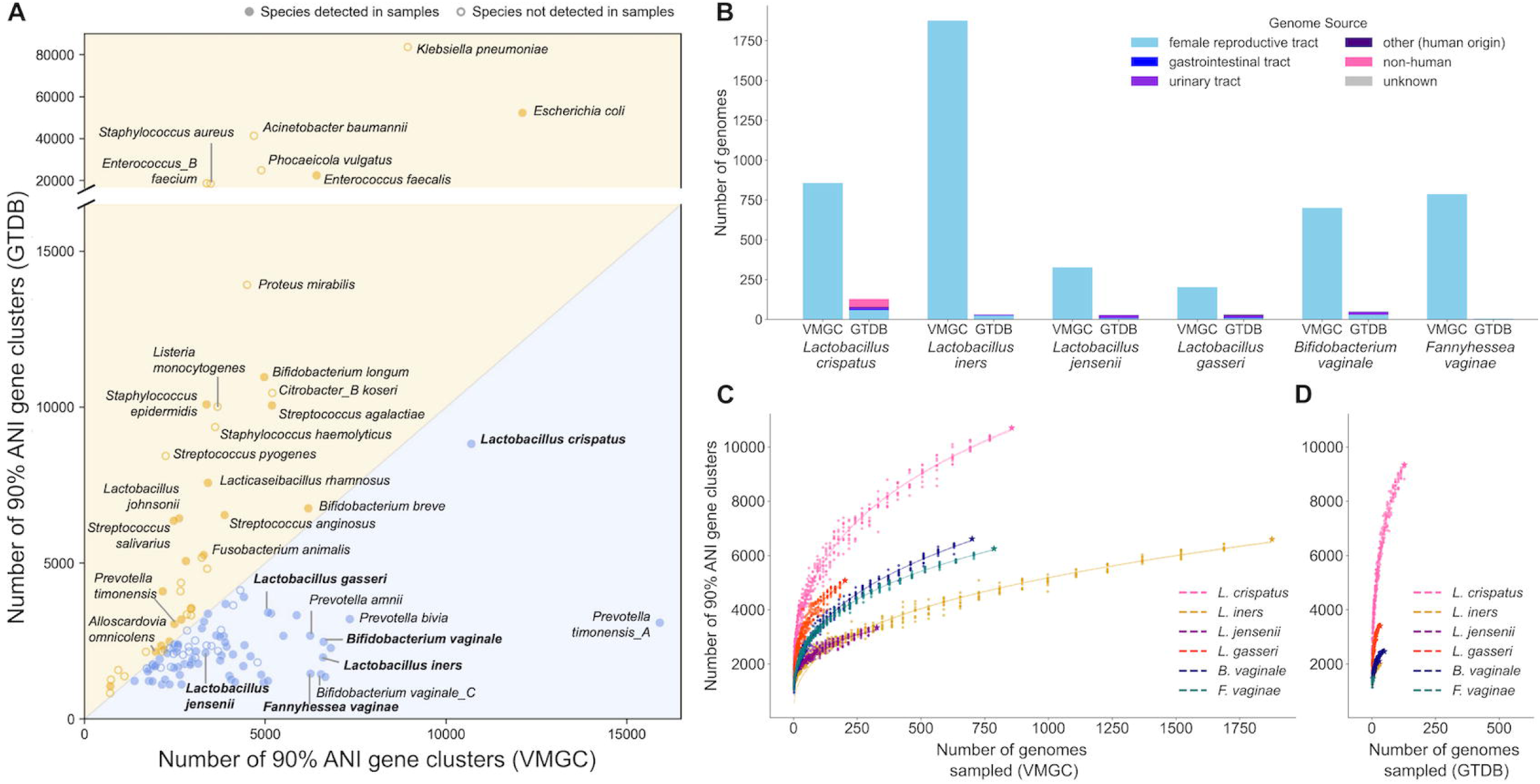
Comparison of pangenome databases constructed from VMGC and GTDB genomes. **A.** Comparison of the number of 90% ANI gene clusters in the pangenome among shared species in databases constructed from VMGC and GTDB genomes. Species with relative abundance greater than 5% in at least 5% of samples are labeled in bold. Species detected in at least one of 352 samples from Norenhag, et al. (2024) are depicted with filled circles, and species not detected in any of these samples are depicted with an unfilled circle. **B.** Comparison of the number of genomes and their origins for six prevalent vaginal microbiome species. *Bifidobacterium vaginale* is a member of the clade referred to as *Gardnerella vaginalis* in NCBI taxonomy, and *Fannyhessea vaginae* was formerly known as *Atopobium vaginae*. See Figure S4 for proportional representation of genome sources by database. **C-D.** Rarefaction of pangenome size for six selected species using subsampled genomes from VMGC (C) and GTDB (D). The number of sampled genomes was decreased by 10% at each subsampling point, with ten replicates at each subsampling point. The generated data points (number of sampled genomes vs. pangenome size) were fit to power functions.

We used MIDAS to quantify the relative abundance of 129 species with one-to-one mappings across GTDB versions among 352 vaginal samples from Norenhag et al. (2024)^29^ (Figure S3). Of the 86 species with relative abundance greater than 1% in at least one vaginal sample, 76% had larger pangenomes in the database built from VMGC genomes. VMGC has a larger pangenome and more input genomes for all six species with relative abundance greater than 5% in at least 5% of samples (Figure 2A-B, S4). We compared pangenome completeness between databases using rarefaction analysis on these six species. Tail slopes from collector’s curves (calculated using the four largest subsampling points per database) were lower in VMGC than GTDB for these six species (Wilcoxon *p* = 0.0312), indicating that VMGC-derived pangenomes are approaching saturation while GTDB pangenomes continue to accumulate novel gene content as more genomes are added (Figure 2C-D). When comparing pangenome sizes at matched genome counts, GTDB and VMGC were similar across species, except for *L. crispatus*, where the GTDB-based pangenome was larger at lower genome counts. The same trend persisted after stratifying *L. crispatus* genomes from the female reproductive tract by genome type (i.e., MAGs vs. isolates) (Figure S5A-B), indicating the discrepancy may be driven by incompleteness in *L. crispatus* MAGs. In *L. crispatus* and other species, isolate genomes from the female reproductive tract harbored a higher number of predicted mobile elements and viral sequences than female reproductive tract MAGs (Figure S5C-D). Although MAGs are generally less complete than isolate genomes, the substantial quantity of vaginal MAGs in the VMGC database offsets their lower per-genome contribution, resulting in a larger total pangenome across all six species.

### VMGC-based pangenome database expands accessory gene repertoire in prevalent vaginal bacteria

Pangenome expansion is most meaningful when it uncovers biologically relevant genes, rather than merely increasing the number of genes associated with a species. To further characterize differences between pangenome reference databases constructed from VMGC and GTDB genomes, we analyzed the overlap and distinctiveness of the gene clusters across databases. We rebuilt pangenome databases with the set of all genomes in VMGC and GTDB for the six prevalent species identified above. The resulting 90% ANI gene clusters were classified as follows: VMGC-only (containing sequences from genomes found exclusively in VMGC), GTDB-only (containing sequences from genomes found exclusively in GTDB), or shared (containing sequences from genomes found in VMGC and GTDB; i.e., at minimum, a single sequence from a genome found in both GTDB and VMGC, or two sequences: one from a genome found only in VMGC and one from a genome found only in GTDB). The number of VMGC-only genes was larger than the number of GTDB-only genes for all six species (Figure 3A).

**Figure 3.**
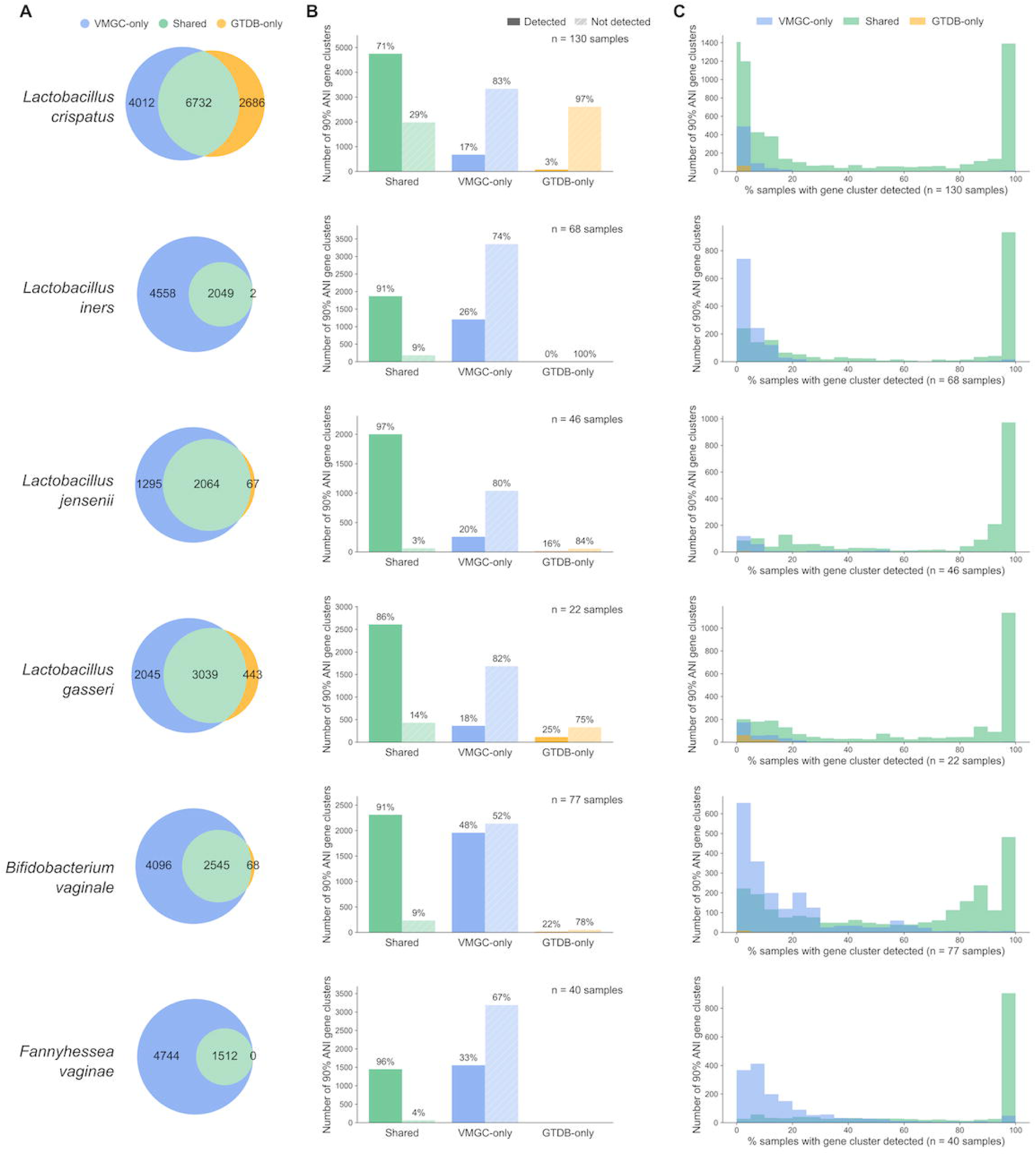
Detection of shared and unique gene clusters in pangenomes constructed from VMGC and GTDB genomes. Each row corresponds to one of six prevalent vaginal microbiome species (labeled at left). **A**. Venn diagrams showing overlap between 90% ANI gene clusters in VMGC and GTDB pangenomes. The number of gene clusters within each group are overlaid on the circles. **B.** Proportion of gene clusters detected (darker color) or not detected (lighter color) in samples from Norenhag et al. (2024), stratified by cluster classification. **C.** Frequency of detection of gene clusters detected in at least one sample, stratified by cluster classification.

To see how commonly the database-specific genes are detected in vaginal samples, we used MIDAS and the combined reference database to measure gene presence and absence in vaginal samples from Norenhag et al. (2024).^29^ While *L. crispatus* has a relatively large number of GTDB-only genes, less than 3% (74/2686) of these genes are detected among the 130 vaginal samples with *L. crispatus* present (Figure 3B). Across all six species, shared genes are more frequently detected than VMGC-only and GTDB-only genes, and more VMGC-only genes are detected than GTDB-only genes (Figure 3B-C). Among genes detected in at least one sample, VMGC-only genes are found at a significantly higher frequency than GTDB-only genes in *L. crispatus*, *L. gasseri,* and *B. vaginale* (two-sided Mann Whitney U *p* = 7x10^-6^, 0.029, and 0.005 respectively). Higher-frequency accessory genes are particularly valuable in association analyses, as they provide greater statistical power to detect associations with clinical traits.

Accessory genes often encode functions that are relevant to fitness in a specific environment. We used a permutation analysis to identify differentially prevalent PFAM domains in VMGC-only and GTDB-only gene clusters. *L. crispatus* GTDB-only centroids were enriched for eleven PFAM domains, including glycosyltransferase domains, cell surface-related domains, and a bacteriophage abortive infection AbiH domain (Table 1). In contrast, *L. crispatus* VMGC-only centroids were enriched for three domains associated with integrated phage sequences: BRO family N-terminal domain, AAA domain, and Terminase_1 (Table 1). *L. crispatus* VMGC-only genomes had a higher number of predicted viral regions compared to GTDB-only genomes (Mann Whitney U *p* = 0.001, Figure 4A). More broadly, *L. crispatus* genomes from human hosts had a higher number of viral regions than those from non-human hosts (*p* = 1x10^-5^, Figure 4B), and genomes from the human female reproductive tract had more predicted viral regions than genomes from the human gastrointestinal tract (*p* = 0.012) and the combined set of genomes from all other human body sites (*p* = 0.014) (Figure 4C). These trends remained consistent when subsetting to only high-quality genomes or only isolate genomes (Figure S6). These findings align with the well-documented abundance of bacteriophages in the vaginal microbiome.^30,31^ Furthermore, they demonstrate that the site-specific database can capture genomic virus-bacteria interactions at the genomic level, even when accounting for the potential limitations of binning-based assembly^32^ and the lower quality of VMGC MAGs relative to isolate genomes.

**Figure 4.**
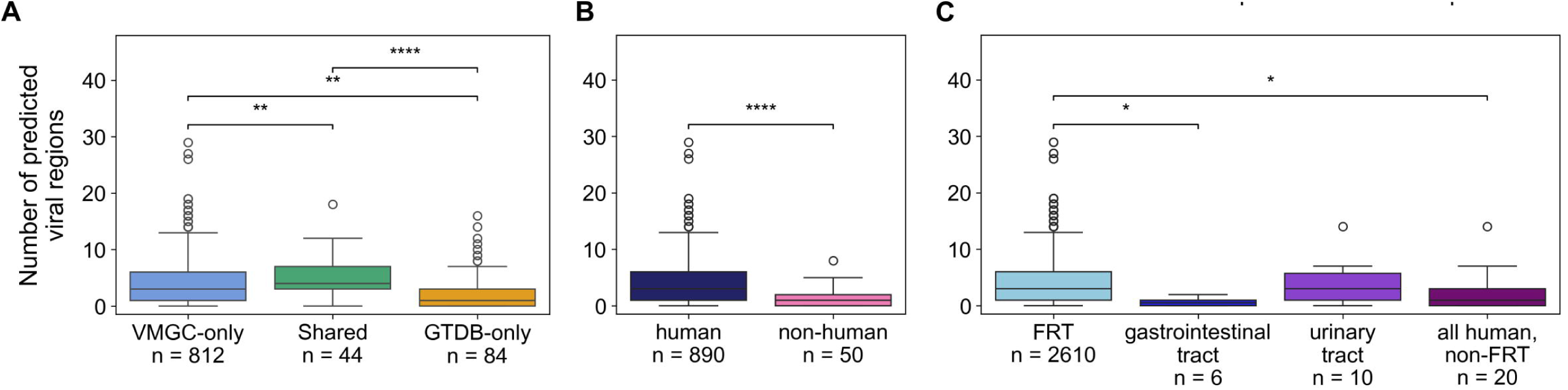
Comparison of number of predicted viral regions among *L. crispatus* genomes, stratified by database category (A), host (B), and human body site (C). Viral regions in genome assemblies were predicted with geNomad. The number of genomes in each group is indicated on the x-axes. Pairwise comparisons within plots were made via a two-sided Mann Whitney U test; pairs with significant differences in number of viral regions are indicated with brackets. **** *p* ≤ 0.0001; *** *p* ≤ 0.001; ** *p* ≤ 0.01; * *p* ≤ 0.05. FRT = female reproductive tract.

**Table 1:** PFAM enrichment.

| Species | Enrichment | PFAM | BH-adjusted *p* |
| --- | --- | --- | --- |
| *Lactobacillus crispatus* | Enriched in GTDB-only centroids | Bacteriophage abortive infection AbiH | 0.0080 |
|  |  | YSIRK type signal peptide | 0.0080 |
|  |  | LPXTG cell wall anchor motif | 0.0080 |
|  |  | Methylase_S | 0.0080 |
|  |  | Glycosyl transferase family 2 | 0.0080 |
|  |  | Glycosyl transferases group 1 | 0.0080 |
|  |  | N-6 DNA Methylase | 0.0140 |
|  |  | Polysaccharide biosynthesis C-terminal domain | 0.0224 |
|  |  | Glycosyltransferase Family 4 | 0.0224 |
|  |  | Hexapep | 0.0420 |
|  |  | SLAP (surface layer-associated protein) domain | 0.0440 |
|  | Enriched in VMGC-only centroids | BRO family, N-terminal domain | 0.0080 |
|  |  | AAA domain | 0.0356 |
|  |  | Terminase_1 | 0.0430 |
| *Fannyhessea vaginae* | Enriched in VMGC-only centroids | CHAP domain | 0.0190 |
|  |  | Methylase_S | 0.0190 |

### Thirteen *Lactobacillus crispatus* accessory genes are associated with increased cervical dysplasia risk

We applied the VMGC pangenome database to shotgun metagenomic data from a Swedish cervical dysplasia cohort^29^ to evaluate its effectiveness in detecting intraspecies associations with clinical phenotypes.

To model these associations, we employed the MWAS-based tool microSLAM.^22^ *L. crispatus* was chosen for analysis as the sole species with sufficient genome coverage to meet microSLAM thresholds (>80 samples) for robust gene detection. Of 131 samples with *L. crispatus*, 82 are from healthy controls and 49 are from individuals with cervical dysplasia. We focused our association analysis on 75% ANI gene clusters, representing broader functional gene groups, to increase statistical power by reducing multiple testing. We identified 832 75% ANI gene clusters detected in 15% to 75% of these samples. We used microSLAM to model the associations between the presence/absence vectors of these genes and disease state, also including *L. crispatus* relative abundance, a propensity score calculated from five clinical metadata variables, and population structure as covariates (see Methods). We identified thirteen accessory genes associated with cervical dysplasia (locFDR < 0.2; OR = 2.48-6.19; Figure 5A, Table S3). These genes encode six hypothetical proteins, a protein with a domain of unknown function, three transcriptional regulators, an ion transporter, and both components of the HicAB type II toxin-antitoxin system. All significant genes are more frequently detected in cervical dysplasia samples than in samples from healthy controls (Figure 5B). The ion transporter is a VMGC-only gene. Two groups of genes have similar co-occurrence patterns across samples (Jaccard similarity >= 0.8) (Figure 5C).

**Figure 5.**
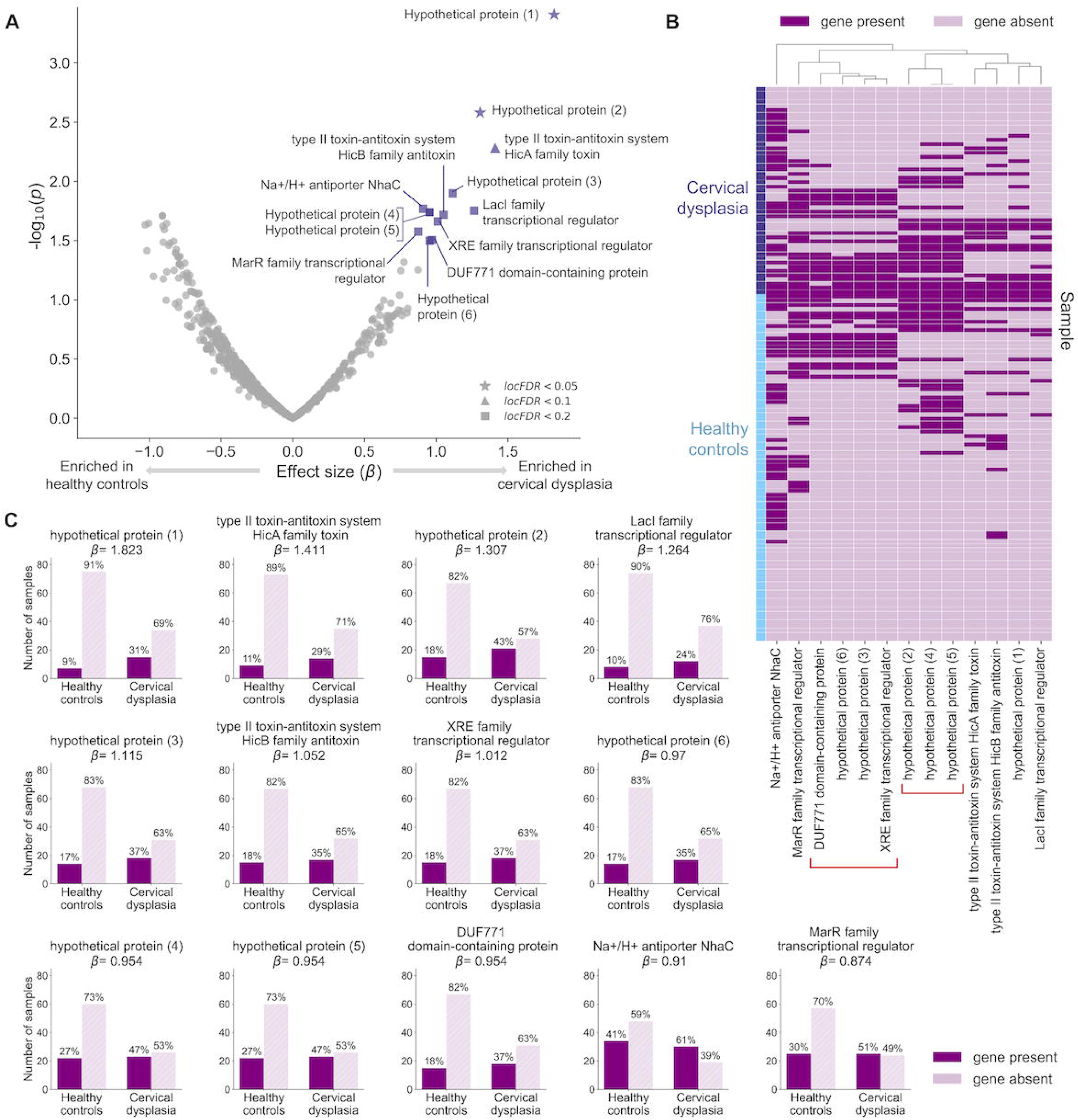
*L. crispatus* accessory genes associated with cervical dysplasia. **A.** Effect size and significance of 832 accessory genes used as input into association analysis. Genes with locFDR < 0.2 are annotated. **B.** Prevalence of significant genes across samples; significant genes are detected in a larger proportion of cervical dysplasia samples compared to healthy controls. **C.** Presence and absence of significant genes across samples, clustered by Jaccard similarity of gene presence/absence vectors. Genes in red brackets have pairwise Jaccard similarity greater than or equal to 0.8 for all genes in the group.

### Genomic context of cervical dysplasia-associated genes

To further elucidate potential functions of the cervical dysplasia-associated genes, we analyzed their genomic context in high-quality *L. crispatus* isolates (Table S4). We identified all *L. crispatus* genome assemblies available on NCBI constructed from single-isolate (non-metagenomic) sequencing and a ‘complete genome’ assembly level. We used BLASTn to identify predicted coding sequences from these 22 strains with homology to the thirteen significant gene sequences. The significant genes are variably detected across the *L. crispatus* isolate genomes (Figure 6A), consistent with our variable detection of them in genomes. The sets of genes with similar co-occurrence patterns across samples were co-located in syntenic regions of the isolate genomes (Figure 6B and 6C). We used Operon-mapper^33^ to predict co-regulated genes in these regions. A group of five cervical dysplasia-associated genes encoding two hypothetical proteins, an XRE family transcriptional regulator, a MarR family transcriptional regulator, and a protein with a domain of unknown function are genomically adjacent, but only the latter two genes are predicted to be co-regulated (Figure 6B). The group of three co-occurring genes encoding hypothetical proteins is predicted to be a part of an operon within an approximately 40 kb region predicted to be a *Caudoviricetes* provirus (Figure 6C, S7). The HicAB toxin-antitoxin system components are predicted to be part of an operon that also includes two hypothetical proteins (Figure 6D). The LacI transcriptional regulator is part of a potential carbohydrate utilization operon (Figure 6E). The remaining hypothetical protein is predicted to be co-regulated with a GNAT family N-acetyltransferase (Figure 6F), and the *nhaC* ion transporter is not predicted to be co-regulated with any other genes (Figure 6G).

**Figure 6.**
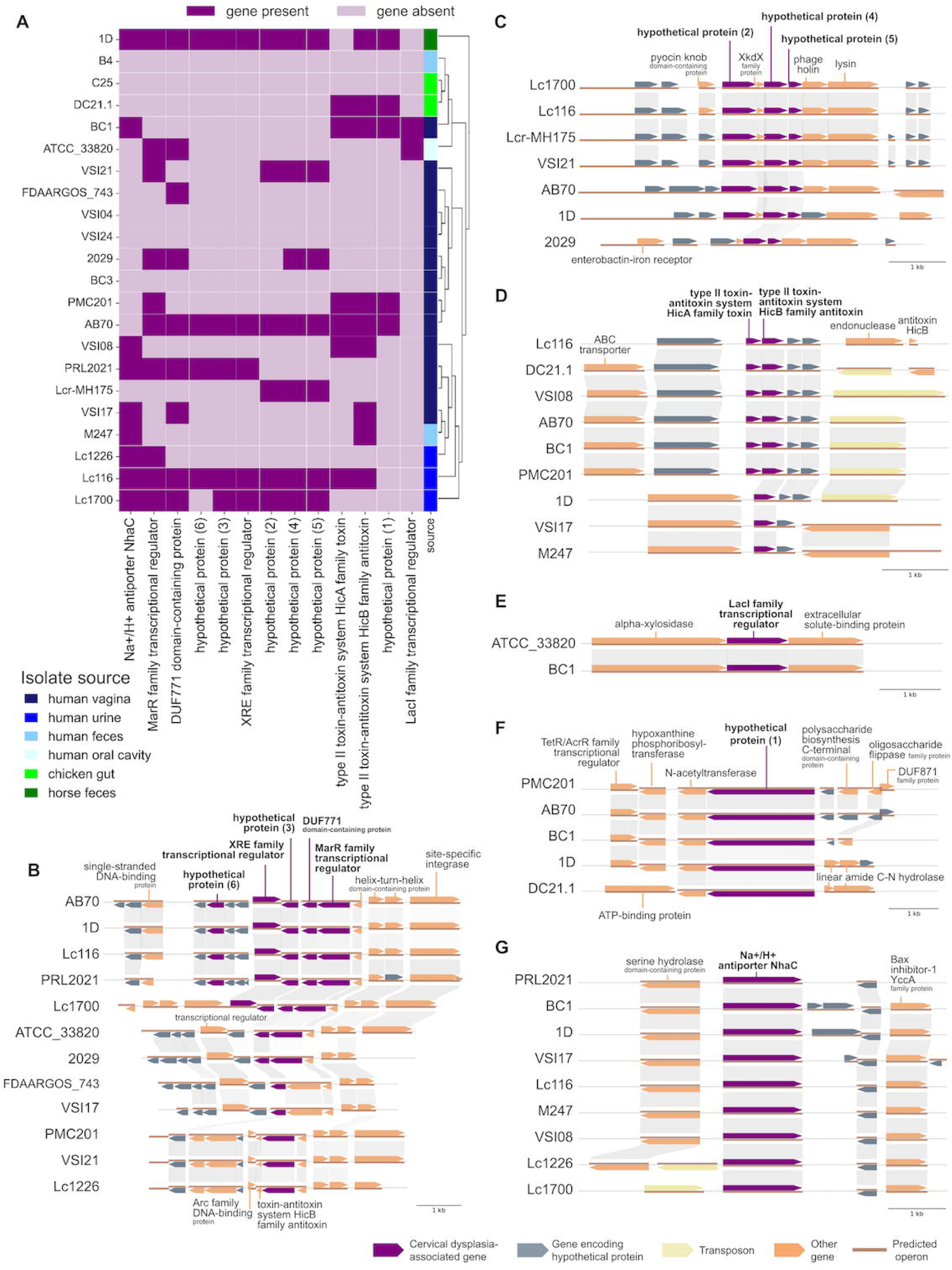
Detection and genomic context of cervical dysplasia-associated genes in NCBI *L. crispatus* isolates. **A.** Presence/absence of cervical dysplasia-associated genes among 22 *L. crispatus* genome assemblies from single-isolate sequencing. The phylogenetic tree on the right side of the plot was constructed via StrainPhlAn using the complete genome assemblies and depicts the branching topology only (branch lengths are arbitrary). The phylogenetic tree with branch lengths scaled to genetic distance is depicted in Figure S7. **B-G.** 5kb genomic context windows around significant genes among NCBI isolates with the given gene(s) detected. Gray connections across isolates indicate genes are reciprocal best hits as identified by MMseqs2, with sequence identity of at least 75% across at least 75% of both sequences. Brown segments indicate operons predicted by Operon-mapper.

To assess the broader phylogenetic distribution of these genes, we used BLASTx^34^ to search for homologous sequences across bacterial species (Table S5). The two top-scoring hypothetical proteins (1 and 2) showed no significant hits outside of *L. crispatus*, suggesting they may be species-specific. Hypothetical proteins 3-6, the XRE family transcriptional regulator, and the DUF771 domain-containing protein had homologs within other members of the *Lactobacillaceae* family. The *nhaC* antiporter, LacI family transcriptional regulator, and MarR family transcriptional regulator were found throughout *Lactobacillaceae* and detected in a limited number of species outside this family. The two toxin-antitoxin system genes showed the broadest distribution, with homologs detected across diverse bacterial species. Collectively, the cervical dysplasia-associated genes span diverse functional categories, genomic contexts, and degrees of taxonomic specificity; the prevalence of proteins with unknown functions in and near the set of cervical dysplasia-associated genes highlights the need for functional studies to investigate the mechanistic basis of these associations.

## DISCUSSION

In this study, we constructed a vaginal microbiome-specific pangenome database using the Vaginal Microbiome Genome Collection and demonstrated its utility in identifying associations between bacterial accessory genes and clinical phenotypes. Previously published MIDAS pangenome databases use genomes largely derived from gastrointestinal and environmental organisms, limiting their ability to reflect the genetic diversity of bacteria from other body sites such as the human vaginal microbiome. The integration of MAGs is especially valuable for expanding pangenomes without the prerequisite of *in vitro* cultivation. This is a critical consideration for the vaginal microbiome, as many resident species are anaerobic and experimentally challenging to culture and isolate.^35^ For hallmark *Lactobacillus* species, as well as dysbiosis-associated organisms like *Fannyhessea vaginae* and *Bifidobacterium vaginale*, use of VMGC genomes increased pangenome size and completeness, with percent increase in size ranging from 21% to 332%. Pangenome expansion uncovered accessory genes that were frequently detected in vaginal samples and enriched for environment-specific functions. For *L. crispatus*, vaginal genomes were enriched for regions of viral origin and PFAM domains associated with integrated phage, consistent with prior observations that viral elements are widely prevalent in the vaginal niche.^31,36^ One of these cervical dysplasia-associated genes (*nhaC* ion transporter) is found in the VMGC pangenome database but not in the GTDB database, highlighting the importance of using a body site-specific database in MWAS analyses.

We identified thirteen *L. crispatus* accessory genes associated with increased risk of cervical dysplasia, despite the species’ overall protectiveness against cervical dysplasia and other adverse reproductive health outcomes.^37^ These include transcriptional regulators, toxin-antitoxin system components, and proteins of unknown function, several of which were embedded within phage-associated regions or putative operons. The HicAB toxin-antitoxin system promotes biofilm formation and inhibits translation to slow bacterial growth in response to stress, promoting persistence during antibiotic treatment and acid stress in *Burkholderia pseudomallei* and *Acetobacter pasteurianus.*^38–40^ Other type II antitoxin systems are involved in abortive infection of phage DNA, and a similar role has been proposed for the HicAB system via comparative genomics.^41^ Given the well-documented relationship between HPV infection and cervical dysplasia,^11^ the significance of three genes of unknown function in a phage-derived region is particularly notable and warrants further study of virus-bacteria interactions in this context. HPV infection and cervical dysplasia are frequently accompanied by dysbiosis, inflammation, pH changes, and barrier disruption; these factors are known to induce stress responses, alter metabolic demands, and drive prophage induction.^42,43^ The significant genes involved in acid stress (*nhaC* ion transporter), sugar metabolism (LacI family transcriptional regulator), and persistence (HicAB toxin-antitoxin system) may promote survival in the hostile, high-inflammation environment associated with cervical dysplasia. Experimental work is needed to understand any mechanistic role of these genes. They may be directly involved in the etiology of cervical dysplasia, act as a response to the environmental changes associated with dysbiosis, or make *L. crispatus* more permissive to the growth of pathogenic species, increasing the likelihood of transitioning to a dysbiotic state that favors viral infection and persistence.

The heterogeneity within *L. crispatus* populations underscores the need to move beyond species-level assessments when studying host-microbiome relationships. In particular, distinguishing beneficial from neutral or potentially pathogenic sublineages may be critical for understanding why *L. crispatus* dominance is not universally protective against dysbiosis and adverse outcomes. Norenhag, et al. 2024^44^ identified amino acid biosynthesis and sugar degradation pathways enriched in healthy controls and nucleotide and peptidoglycan biosynthesis pathways enriched in cases of cervical dysplasia.^29^ *L. crispatus* strongly contributed to the pathways enriched in healthy controls, but within-species variation of these pathways was not reported. This is the first study to our knowledge to analyze bacterial intraspecies variation in relation to cervical dysplasia and points to the analytic gain from a richer, more tailored, reference database.

Unlike existing scripts for constructing a MIDAS-compatible pangenome database, our Nextflow pipeline is automated, highly parallelized, and compatible with multiple executors. This workflow may be particularly useful for constructing pangenome databases from genome collections recently published for the oral and skin microbiomes,^26,27^ as well as keeping reference sets updated and up-to-date with new data. VIRGO2, an expansive catalog of non-redundant vaginal microbiome genes pooled across species, is also an excellent resource for measuring genetic diversity in the vaginal microbiome.^45^ Our reference database contains pangenomes constructed for each species individually, and adjustable ANI clustering thresholds to suit the desired level of granularity. Our approach may be particularly advantageous for studies focused on specific species or for seamless integration with the MIDAS pipeline.

This study has several limitations. We did not stratify cases by severity of dysplasia, as the sample sizes within each subgroup would be too small for statistical analysis. Access to healthcare influences the availability of cervical dysplasia screening and screening practices vary across countries,^46^ so individuals included in this study may not fully represent the broader population of people with and without cervical dysplasia; generalizing these results to other populations will require validation in independent cohorts. Furthermore, users of the VMGC reference should consider several limitations inherent to reference-based approaches. Contributing isolate genomes and MAGs in VMGC are predominantly derived from samples from the United States, China, and Europe,^47^ which may limit the representation of genetic diversity from other geographic regions. The use of MAGs may introduce bias from assembly errors, chimeric contigs, or incomplete gene predictions.^32^ However, these challenges can be mitigated using improved assembly algorithms, and our reproducible Nextflow pipeline allows pangenomes to be easily reconstructed as MAGs are refined. Future users should evaluate pangenome size, saturation, and quality of contributing genomes when deciding whether the pangenome reference for a particular species is sufficiently comprehensive for their application or whether de novo gene content characterization would be more appropriate. Our methodology for accurate measurement of gene content relies on sufficient sequencing depth across at least 80 samples. This technical constraint limited our scope to *L. crispatus*, preventing exploration of potential gene-disease associations in other bacterial species. Moreover, while our model accounts for the relative abundance of *L. crispatus*, it does not incorporate the broader taxonomic composition of the vaginal microbiome. The top dysplasia-associated gene is a hypothetical protein unique to *L. crispatus* with no known protein domains, underscoring the need for approaches to improve the functional annotation of vaginal pangenomes. Finally, the significant genes identified here are associations rather than causal determinants of cervical dysplasia; experimental validation is necessary to confirm their roles and characterize the genes with unknown function.

Together, our work establishes both a resource and a methodological framework for expanding MIDAS pangenome databases to better represent body site-specific microbiomes. The VMGC-based database improves the resolution of metagenomic analyses in the vaginal microbiome, reveals novel accessory genes of potential clinical importance, and demonstrates that intraspecies diversity within key taxa like *L. crispatus* may shape reproductive health outcomes. Extending similar approaches to other body sites and diseases will help uncover overlooked functional variation within the human microbiome and elucidate its role in health and disease.

## METHODS

### Categorization of GTDB genome sources

We downloaded metadata for the 258,405 GTDB genomes used in the GTDB-based pangenome database via the NCBI datasets command line tool version 18.0.2.^48^ We identified 98,680 genomes with the words “human” or “homo sapiens” in the Assembly BioSample Host, Assembly BioSample Source type, Assembly BioSample Isolation source, Assembly BioSample Models, and/or Assembly BioProject Lineage Title. We manually categorized the 187 most common isolation sources into the following categories: skin, gastrointestinal, respiratory, female reproductive tract, oral, blood, urine, other, unknown, and non-human. These manual annotations categorized the source of 91% (89,483/98,680) of human-associated genomes. We used the gemma-2-9b-it^45^ large language model to categorize the 3,022 remaining isolate sources corresponding to 9% of genomes. We removed genomes whose sources remained unknown (n = 33,573) or were recategorized as non-human (n = 532) before calculating the proportions of genomes isolated from each body site. For the six species depicted in Figure 2B and S4, we manually verified and re-annotated genomes whose sources were categorized as unknown and non-human through a literature search.

### Genome assembly download

Prokaryotic vaginal genomes and their taxonomic assignments to GTDB r214.1 were downloaded from Huang et al. (2024).^25^ Accession numbers and GTDB r202 species assignments for 258,405 genomes in the preconstructed pangenome database generated from GTDB r202 genomes were downloaded via MIDAS version 3.0.1.^20^ We used gtdb-taxdump version v0.5.0^50^ to map taxonomic assignments across versions r202 and r214.1 of GTDB. Species not present in both database versions and species with many-to-one mappings across versions were excluded from the comparison of pangenome databases. We verified that genomes from the remaining species were assigned to the same species in GTDB versions r202 and r214.1, and we downloaded GTDB genome assemblies via the NCBI datasets command line tool version 18.0.2.^48^

### Nextflow pipeline for constructing custom pangenome databases

Nextflow is an HPC and workflow orchestration system widely used in biomedical research notable for versatility and scalability. We created a Nextflow pipeline to streamline construction of a pangenome database from a user-specified set of genomes and their species assignments. The steps for constructing a pangenome reference database are described in Zhao et al. (2023)^21^ and Smith et al. (2025)^20^ and Figure 1A. Briefly, each genome is annotated via Prokka,^51^ and predicted protein-coding sequences are clustered at 99% ANI using VSEARCH.^52^ The centroids of each cluster are reclustered at 95, 90, 85, 80, and 75% ANI, and refined using CD-HIT.^53^ Centroid sequences of 99% ANI clusters are functionally annotated using eggNOG.^54^ For each genome, regions associated with antimicrobial resistance, viral elements, and plasmid/mobile element-associated regions are identified using ResFinder,^55^ geNomad,^56^ and MobileElementFinder,^57^ respectively. We verified that our Nextflow pipeline outputs identical results to the original database build scripts published in Smith et al. (2025)^20^ for two sample species. The database build from VMGC genomes took 199 CPU days (16 real-time days) on a high-performance computing cluster, distributed across 506 nodes equipped with Intel and AMD processors and 12-256 CPUs and 48-1512 GiB of RAM per node. Most processes used 4 or fewer CPUs and less than 16GB of RAM; the Nextflow pipeline dynamically increased CPU and RAM requests for more computationally intensive processes. While this work was completed on a Sun Grid Engine-managed cluster, Nextflow enables easy transfer to SLURM, AWS, GCP, and other HPC systems.

### Cervical dysplasia samples

We downloaded shotgun metagenomic sequencing reads from European Nucleotide Archive projects PRJEB72778 and PRJEB72779 and corresponding metadata from Norenhag et al. (2024).^29^ This is an age-matched case-control study conducted in Sweden (N=177 healthy controls and N=177 cervical dysplasia). We used Geneshot^58^ to run a quality control workflow on the sequencing data, using Trim Galore version 0.6.6^59^ to remove adapter sequences, bwa version 0.7.18^60^ to identify sequences mapping to the human genome version hg38, and samtools version 1.20^61^ to filter out the non-microbial reads. We removed samples with fewer than 250,000 reads remaining after quality control, leaving 352 samples for analysis (175 samples from healthy controls and 177 samples from individuals with cervical dysplasia).

We quantified species-level composition in each sample using the MIDAS run_species command and merged results across samples with the merge_species command. We considered a species to be present in a sample if the median coverage across single-copy marker genes was at least 0.75 and relative abundance was at least 1%. 138 and 137 species were detected in at least one sample when using the VMGC and GTDB-based pangenome database as a reference, respectively.

### Rarefaction analysis

We selected six species with a relative abundance greater than 5% in at least 5% of samples for rarefaction analysis. For each species and database, rarefaction was performed by selecting a subset of genomes, rebuilding the pangenome database with only those genomes as input, and calculating the number of 90% average nucleotide identity gene clusters in the resulting pangenome database. The number of genomes selected began with n-1 genomes, where n is the total number of genomes available for that species, and decreased by 10% until 2 genomes remained. Ten replicates at each number of genomes were performed.

### Gene cluster uniqueness

Genomes from the six species selected above were assigned categories: VMGC-unique (only present in the VMGC database), GTDB-unique (only present in the GTDB database), or common (present in both the VMGC and GTDB databases). For each species, we rebuilt the pangenome database with the combined set of VMGC-unique, GTDB-unique, and common genomes as input. 90% ANI gene clusters were categorized as “VMGC-only” if they only contained sequences derived from VMGC-unique genomes, “GTDB-only” if they only contained sequences derived from GTDB-unique genomes, and shared if they contained sequences from a common genome or sequences from both VMGC-unique and GTDB-unique genomes.

### PFAM enrichment

We used eggNOG-generated PFAM domain annotations to test for functional enrichment in VMGC- and GTDB-only gene clusters for each species. N_clusters_ represents the total number of VMGC-only or GTDB-only 90% ANI gene clusters for a species, and N_PFAM_ represents the number of database-unique gene clusters with a given PFAM domain. For each PFAM with N_PFAM_ greater than or equal to ten, we took 10,000 samples of N_clusters_ gene clusters from the set of all gene clusters for the species. We used the proportion of samples with more than _NPFAM_ gene clusters with the given PFAM annotation as an empirical *p* value. *p* values for each species were adjusted for multiple hypothesis testing via the Benjamini Hochberg procedure.^62^

### Gene presence/absence

The MIDAS run_genes command was used to measure gene content in each vaginal microbiome sample for all species detected in the sample with median coverage of marker genes greater than 2 and reads mapping to at least 50% of marker genes. Gene clusters with copy number of 0.35 or greater were considered present in a sample. We observed that gene detection varied significantly at low sequencing depths. To ensure reliable gene presence/absence determination, we identified the minimum mean depth required for gene counts to stabilize for each species. For most species (*L. crispatus, L. jensenii, L. gasseri, F. vaginae*, and *B. vaginale*), we set the threshold at 5X, where the correlation between depth and gene count was not significant (Spearman *p* > 0.05). For *L. iners*, we iteratively increased the cutoff and determined that a higher threshold of 8X was necessary to eliminate this significant correlation (at 5X coverage Spearman *p* = 0.0005; at 8X coverage Spearman *p* = 0.21). Results of the gene-level analysis were merged across samples via the merge_genes command, with genes clustered at 90% ANI for pangenome database comparisons and 75% ANI for the cervical dysplasia association analysis. The six-species combined database was used as a reference for the database comparisons (Figure 3-4) and the pangenome database constructed from VMGC genomes was used as a reference in the cervical dysplasia association analysis (Figures 5-6).

### Propensity score calculation

For each of 21 metadata variables reported in Norenhag et al. (2024), we constructed a logistic regression model: outcome ∼ variable + 1, with outcome = 0 representing healthy controls and outcome = 1 representing cervical dysplasia. Five variables with *p* < 0.1 in the univariate models were used in the propensity score: LowRisk_HPV, PotentialHR_HPV, Gyn_problem_24hrs, Menses_yesno, and Gyn_BloodyDischarge. Each of these variables is binary. LowRisk_HPV and PotentialHR_HPV did not have missing values for any samples. The variable Menses_yesno was missing data for 7% of samples; we imputed missing values for this column based on subject age. Subjects with AgeAtSampling < 52 were imputed with True, and subjects with AgeAtSampling >= 52 were imputed with False. 52 was selected as the threshold because the average age of menopause is 51-52.^63^ Gyn_problem_24hrs and Gyn_BloodyDischarge had low frequency of nonzero values (16% and 2% of samples, respectively), and for these columns we imputed missing values (10% of samples for both variables) with the mode (0). We fit a logistic regression model with the matrix of selected variables as features and outcome as the response variable. The predicted probability of each sample belonging to the disease group based on this model was used as a propensity score.

### Cervical dysplasia association analysis

We used the microSLAM R package version 1.0.0.9^22^ to model associations between *L. crispatus* gene content and outcome (*y*). We calculated fixed effects of covariates (α) using a baseline generalized linear model (GLM): y ∼ propensity score + *L. crispatus* relative abundance. We constructed a gene presence/absence matrix from 75% ANI gene clusters detected in 15-75% of samples with mean depth of at least 5 across detected *L. crispatus* genes (n = 702 genes and 131 samples). Next, we calculated a gene relatedness matrix (GRM), a pairwise similarity matrix based on gene presence/absence data used as a proxy for population structure across samples. To calculate the random effects between population structure and outcome (*b*), we combined the baseline GLM and the GRM in a random-effects generalized linear model. Permutation of the tau variable from this model indicated there was not a significant correlation between population structure and outcome (*p* = 0.34). Lastly, we fit a mixed effects model for each gene presence/absence vector *g:*

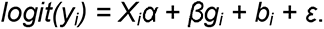

where *i* represents a sample, *X* is the covariate matrix, and β represents the effect size of the gene. All models and statistical tests except for the baseline GLM were implemented via microSLAM. We used the locfdr R package version 1.1.8^64^ to calculate false discovery rates from the *z* scores derived from the mixed effects model. Significant *L. crispatus* gene clusters were annotated via BLASTx^34^ of the centroid sequences against the ClusteredNR protein database. To determine if these genes were present in other species, we filtered for homologs meeting UniRef50 criteria (>50% identity, >80% query coverage) while excluding matches to *L. crispatus*.

### Analysis of *L. crispatus* isolate genome assemblies

We identified 22 *L. crispatus* genome assemblies available on NCBI with sequencing data from a single isolate, assembly level of “complete assembly,” and available coding sequence annotations. Accession numbers for these isolates are provided in Table S4. We used BLASTn version 2.16.0^34^ with minimum query and subject coverage of 75% and minimum identity of 75% to identify predicted coding sequences in each strain with homology to the centroids of 75% ANI gene clusters for *L. crispatus*. The gene with the highest bit score was selected for each coding sequence. If a gene did not have any matches from a strain, it was considered absent in that strain. We used the Operon-mapper web tool^33^ to predict the location of operons and geNomad^56^ to predict viral regions in each strain. PyGenomeViz version 1.6.1^65^ was used for visualizations in Figures 6B-G and S8. The PyGenomeViz integration of MMseqs2 reciprocal best hit search identified sequences with identity of at least 50% across at least 80% of the query and target sequences. The phylogenetic tree for these isolates was created using StrainPhlAn version 3.1.68,^66,67^ with parameters --diversity low, --trim greedy, and –accurate. The closely related species *Lactobacillus helveticus* was selected as the outgroup^68^, and the *L. helveticus* reference strain TCI357 genome assembly was downloaded from NCBI (Table S4). The phylogenetic tree was visualized with ITOL.^69^ The full phylogenetic tree scaled by branch length is available in Figure S7.

## Data Availability

The VMGC pangenome database is publicly available on Zenodo (DOI: 10.5281/zenodo.18355814), with genes clustered at 75%, 80%, 85%, 90%, 95%, and 99% average nucleotide identity for each of 179 species. The Nextflow pipeline for constructing custom MIDAS-compatible pangenome databases is available at https://github.com/clairedubin/MIDASv3_database_build_nf and code for data analysis presented here is available at https://github.com/clairedubin/pangenome_cervical_dysplasia_analysis. The GTDB-based pangenome database was published in Zhao, et al. 2023^21^ and can be downloaded through the MIDAS package.

## STORMS Checklist

The STORMS checklist for this project is available at: https://github.com/clairedubin/pangenome_cervical_dysplasia_analysis/blob/main/STORMS_Excel_1.03.xlsx

