## Supplementary Figures 1-8 for "Expanding vaginal microbiome pangenomes via a custom MIDAS database reveals *Lactobacillus crispatus* accessory genes associated with cervical dysplasia"

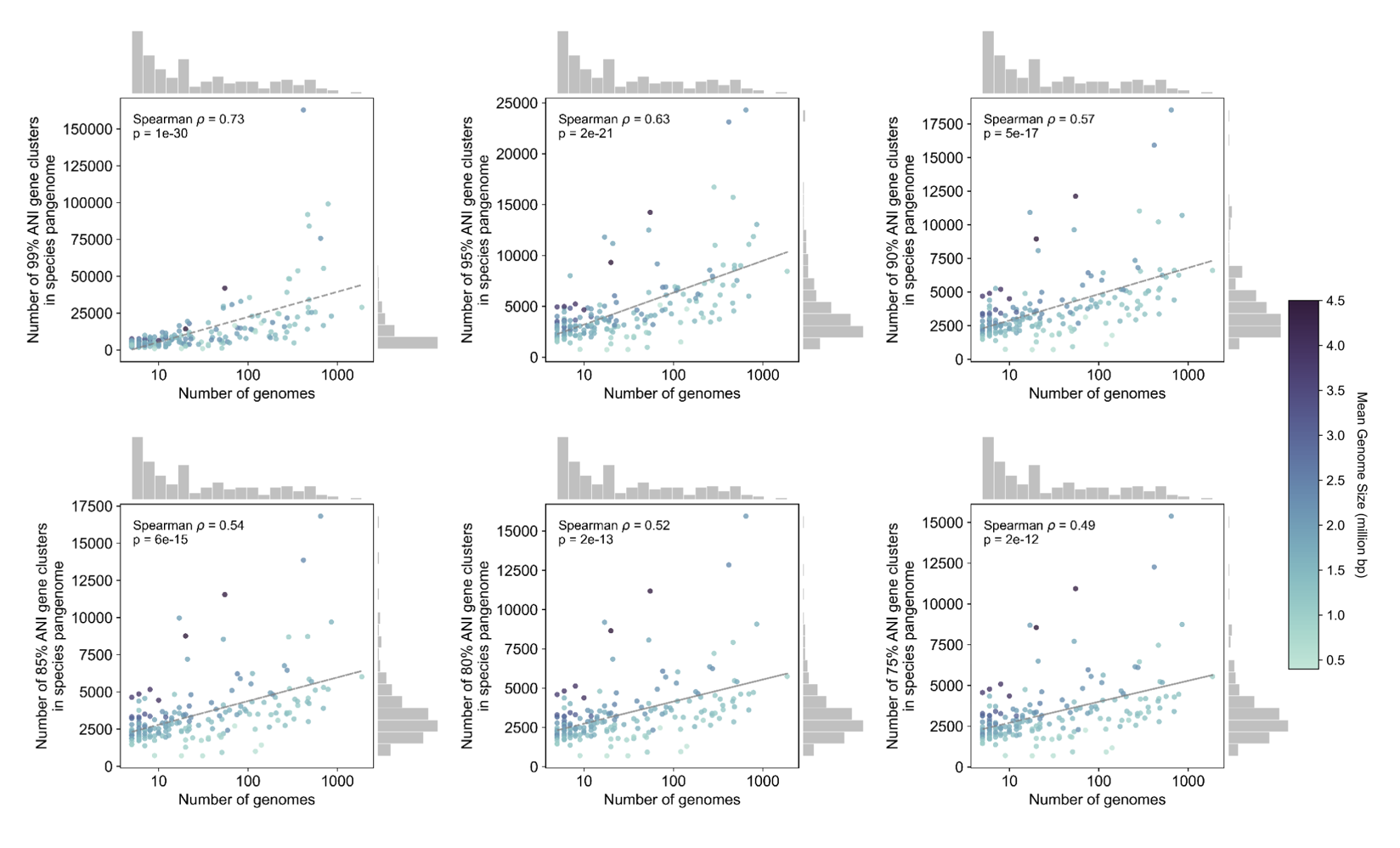

**Figure S1.** **Comparison of number of genomes and VMGC pangenome size with genes clustered at various ANI thresholds.** Species with at least 5 genomes in VMGC are included and ANI thresholds are indicated on the y-axis labels. Each point represents a species, and points are colored by average genome size for the species. Marginal histograms show the distributions of each variable: number of genomes per species (top) and number of gene clusters in species pangenome (right).

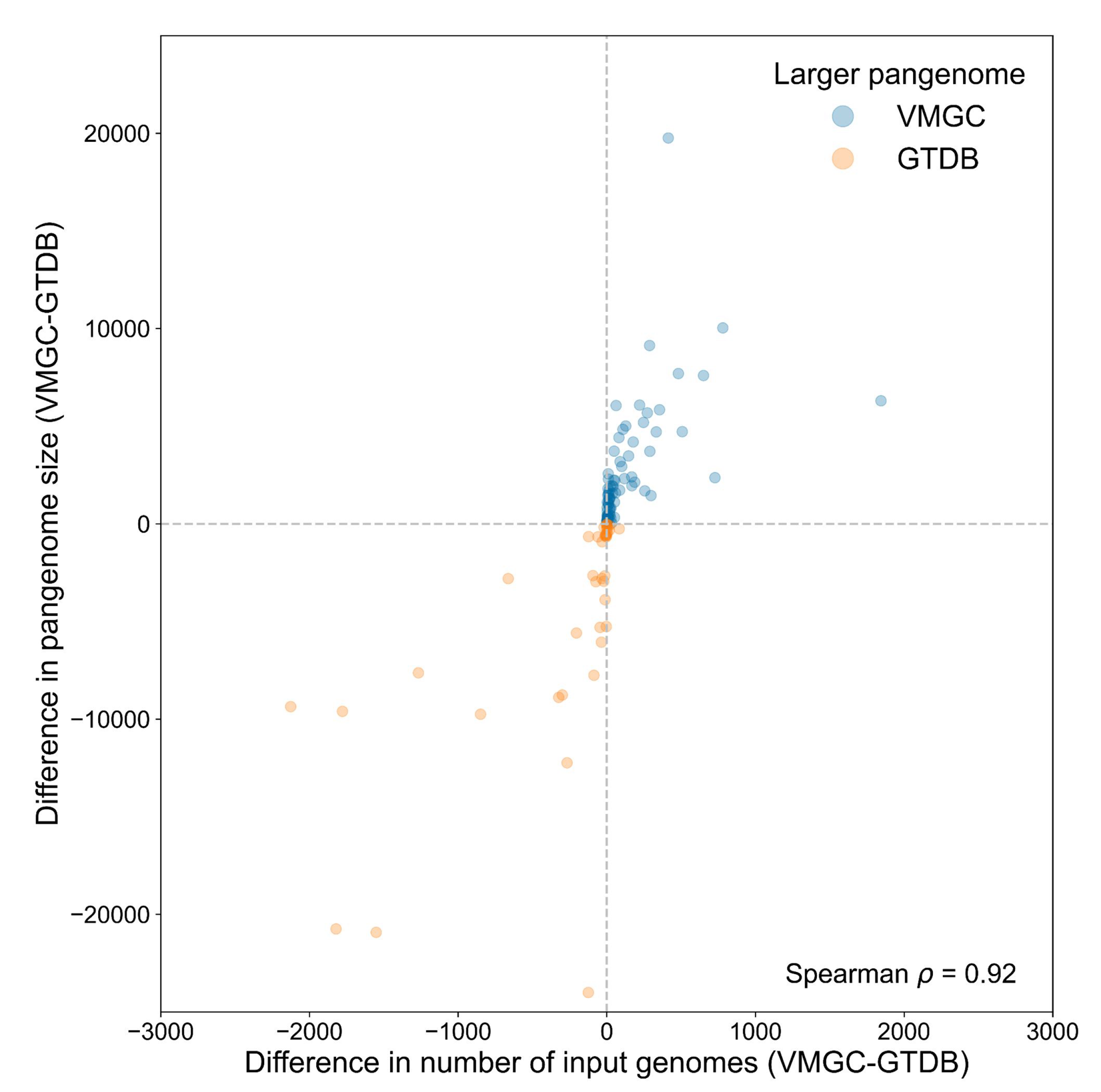

**Figure S2. Difference in pangenome size is positively correlated with differences in number of input genomes.** Each point represents a species. Four species (*Escherichia coli, Klebsiella pneumoniae, Staphylococcus aureus, Acinetobacter baumannii*) with a difference in number of genomes less than -4,000 and a difference in pangenome size less than -20,000 are excluded for clarity.

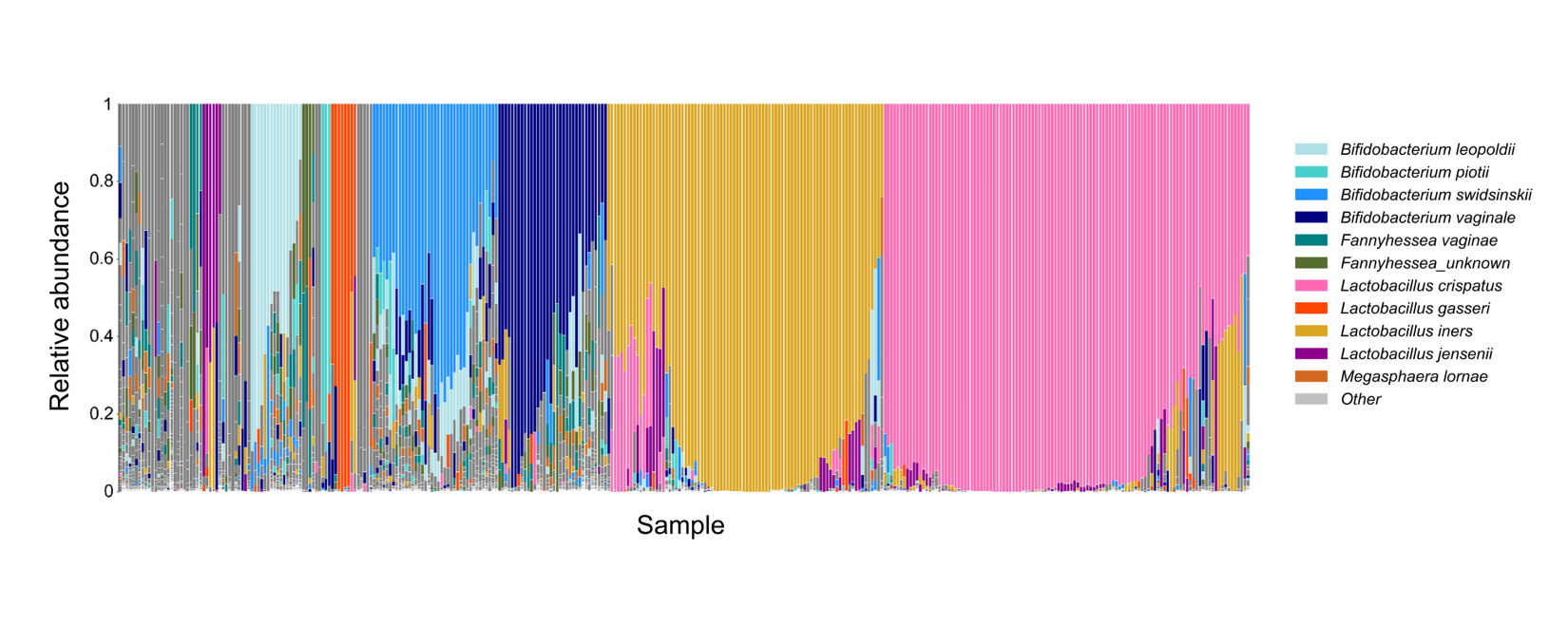

**Figure S3. Relative abundance of prokaryotic species across 352 vaginal samples (from healthy controls and people with cervical dysplasia) from Norenhag et al. (2024).** Relative abundances were calculated via the MIDAS run_species command, using markers from the VMGC-based pangenome reference database. Species with a relative abundance of 5% or more in at least 5% of samples are shown in color and listed in the legend; lower abundance/prevalence species are grouped in gray. *L. crispatus* (pink) is the most abundant species across samples, followed by *L. iners* (gold).

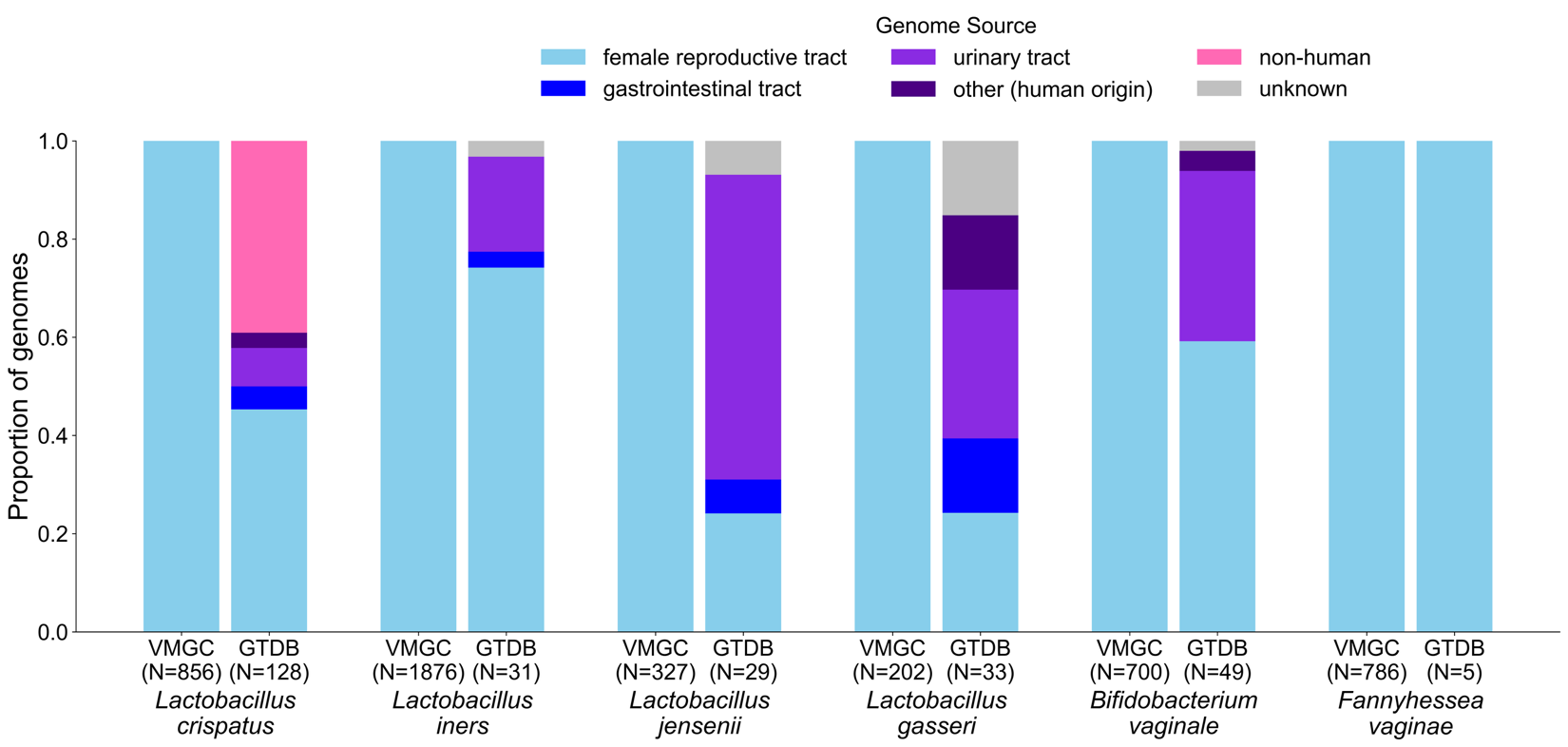

**Figure S4. Proportional representation of genome sources in VMGC and GTDB databases.** Data are as in Figure 2B but displayed as proportions rather than raw counts. The number of genomes for the given species and database are listed below each bar.

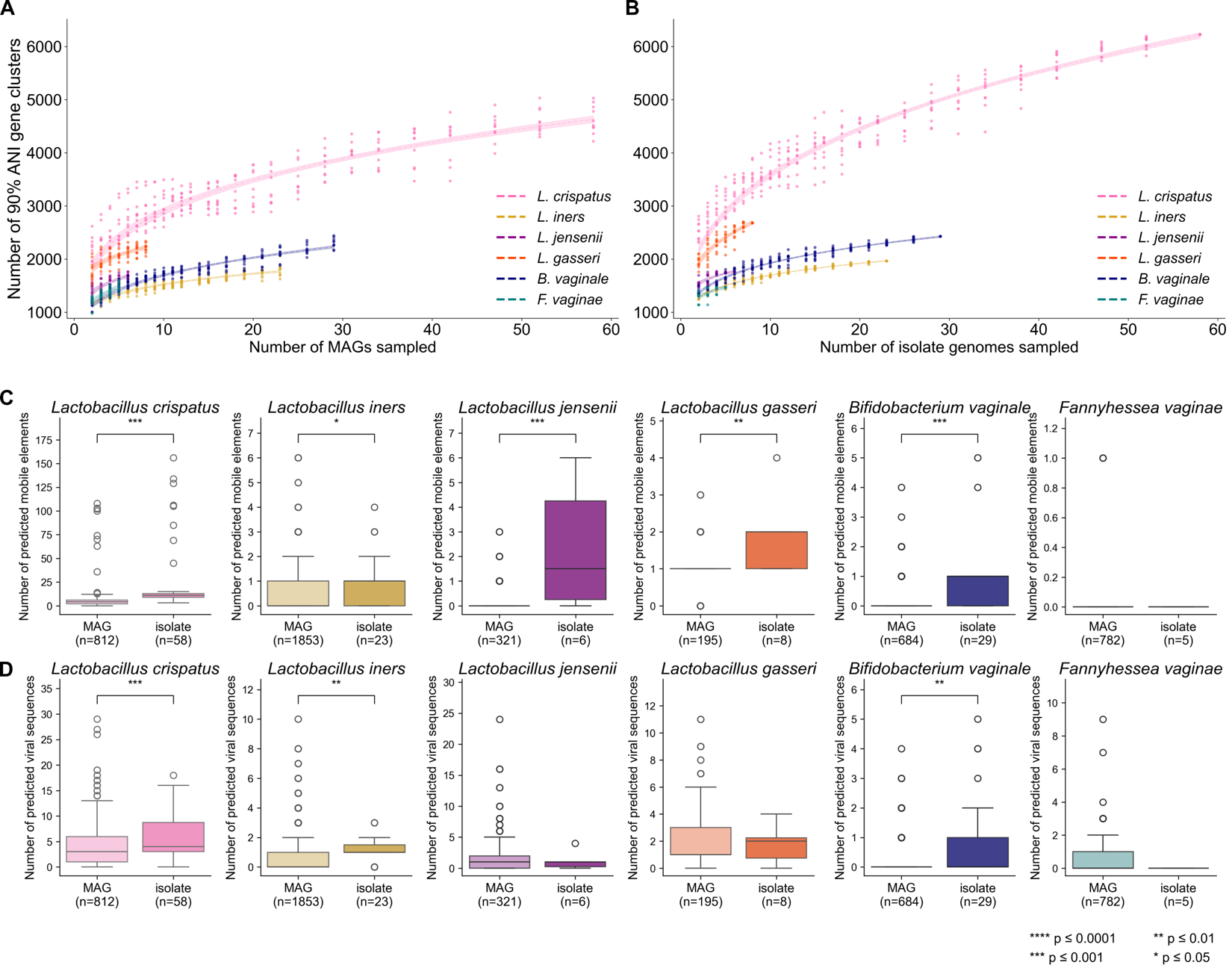

**Figure S5. Comparison of MAGs vs. isolate genomes from the female reproductive tract (FRT). A-B.** Rarefaction of pangenome size among MAGs (**A**) and isolate genomes (**B**). The number of sampled genomes was decreased by 10% at each subsampling point, with ten replicates at each subsampling point. The generated data points (number of sampled genomes vs. pangenome size) were fit to power functions. Resulting pangenome sizes were similar across all species except *L. crispatus*, which had a larger pangenome when using isolate genomes compared to MAGs at matched subsampling points. **C-D.** Comparison of the number of predicted mobile genetic elements (**C**) and viral sequences (**D**) among MAGs and isolate genomes. Mobile genetic elements were predicted by MobileElementFinder and include transposons, integrative and conjugative elements, and insertion sequences. Viral sequences were predicted by geNomad. All data are limited to genomes from the female reproductive tract to avoid confounding with body site. Pairwise comparisons in (**C**) and (**D**) were conducted using the Mann Whitney U test; pairs with significant differences in number of predicted mobile genetics elements or viral regions are indicated with brackets. **** *p* ≤ 0.0001; *** *p* ≤ 0.001; ** *p* ≤ 0.01; * *p* ≤ 0.05.

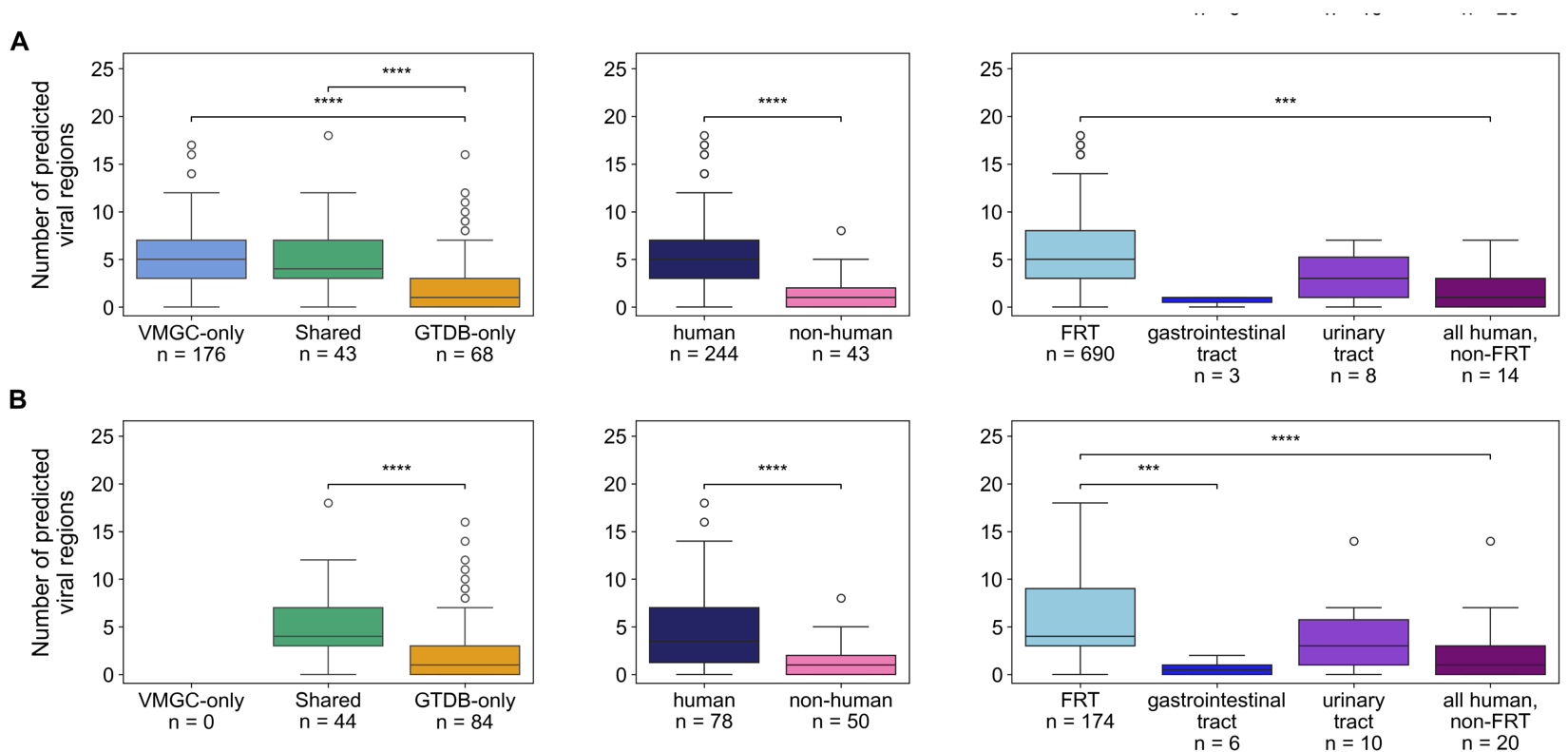

**Figure S6. Comparison of number of geNomad-predicted viral regions among *L. crispatus* genomes, stratified by database category, host, and human body site.** Data are as in Figure 4, but **(A)** is limited to high-quality genomes (completeness > 90% and contamination < 5%) and **(B)** is limited to isolate genomes. Pairwise comparisons within plots were made via a two-sided Mann Whitney U test for all categories with at least 5 genomes; pairs with significant differences in number of viral regions are indicated with brackets. **** *p* ≤ 0.0001; *** *p* ≤ 0.001; ** *p* ≤ 0.01; * *p* ≤ 0.05. FRT = female reproductive tract.

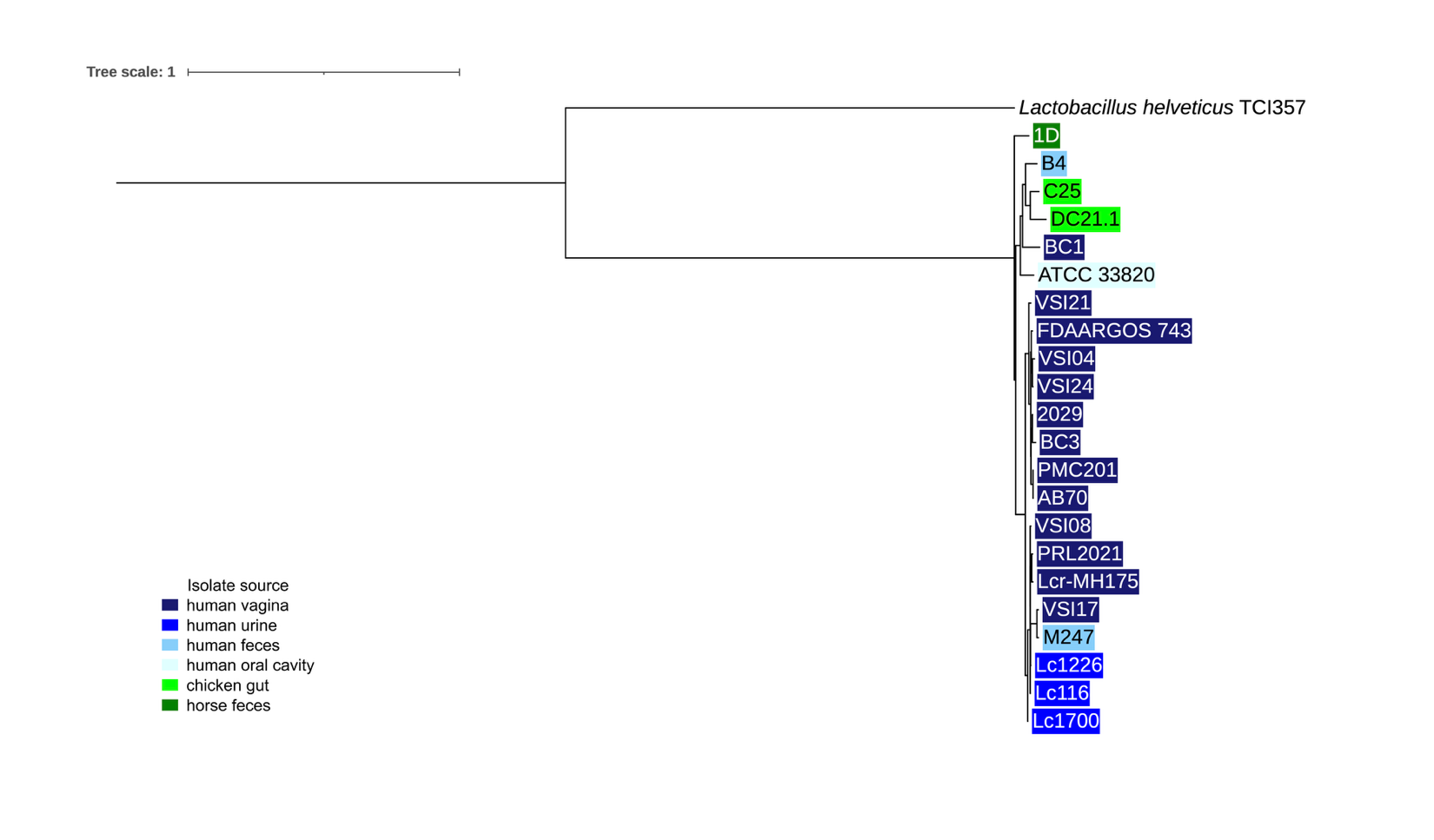
 **Figure S7.** **Phylogenetic tree of *L. crispatus* isolates, with *L. helveticus* strain TCI357 as an outgroup.**

The phylogeny was constructed via StrainPhlAn and visualized with ITOL, rooted at the tree midpoint and scaled by branch length. Branch labels for *L. crispatus* isolates are colored by isolate source.

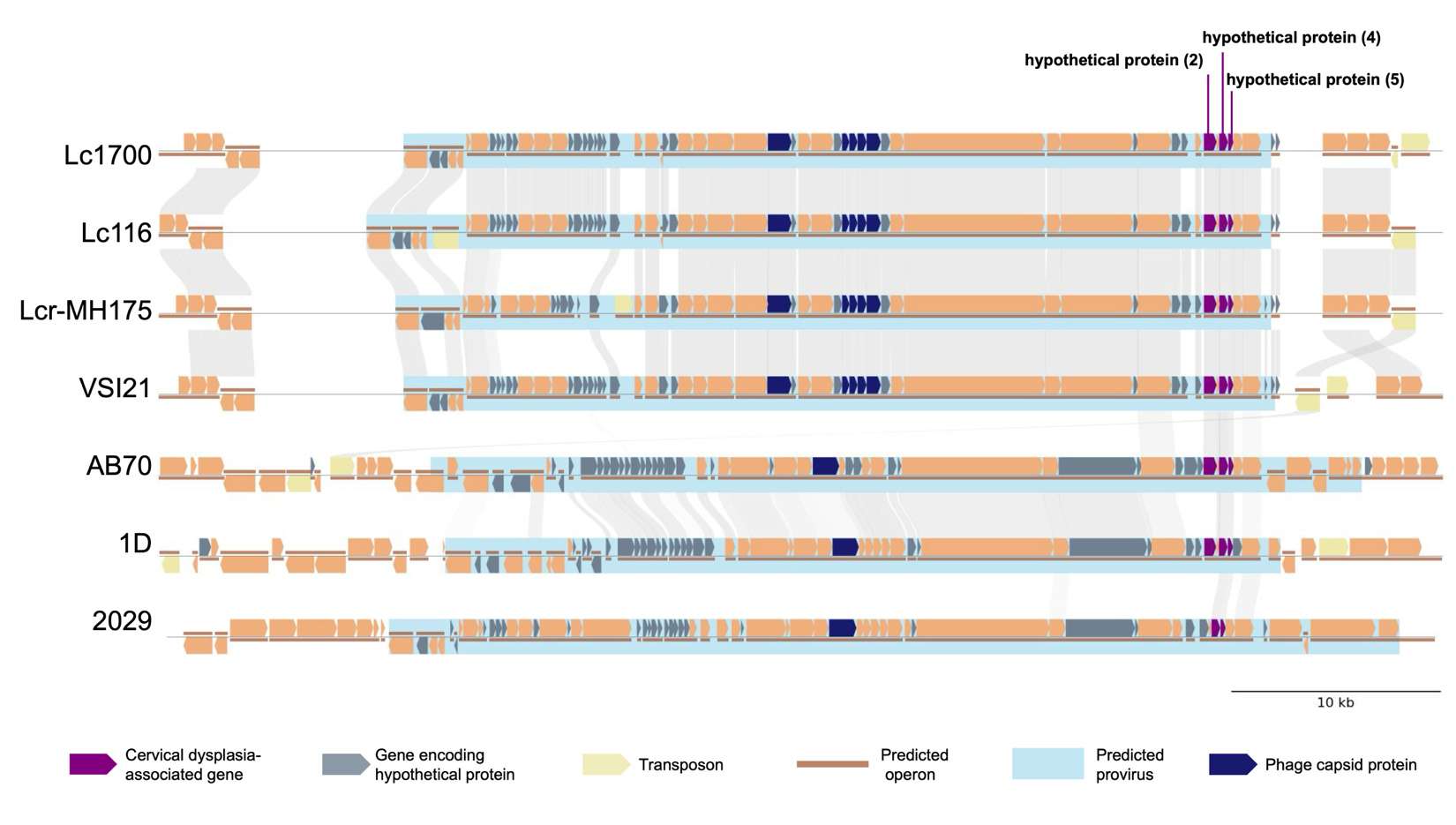

**Figure S8. Three cervical dysplasia-associated hypothetical proteins are located within a 40 kb *Caudoviricetes* provirus region.** Data are as in Figure 6C, with a wider genomic window and showing the predicted provirus region (predicted by geNomad) and phage capsid proteins in light blue and dark blue, respectively.
